# The relationship between interpregnancy interval (IPI) and mother’s preceding pregnancies (PP)

**DOI:** 10.1101/022111

**Authors:** Michel C. Vernier, Michael Schulzer, Pierre R. Vernier

## Abstract

**Introduction:** Following the investigation of the mother’s preceding pregnancies on fetal development and postnatal survival of the neonate, we turned our attention to an earlier period, that is the interval separating the onset of the current pregnancy from the end of the preceding one. The objectives of this study is to investigate the variations of interpregnancy interval length associated to the mother’s preceding pregnancies.

**Methods:** A population of 7773 neonates, alive at the time of hospital discharge, were divided into cohorts according to the current neonate’s sex and number and sex of the mother’s preceding pregnancies. Interpregnancy interval average of each cohort of same neonate’s sex and mother’s parity, but different configuration of preceding pregnancies, were measured and compared.

**Results:** A positive association was found between mother’s preceding pregnancies and length of interpregnancy interval when current pregnancy and preceding pregnancy were of the same sex, and a negative association when they were of opposite sex.

**Discussion:** Interpregnancy interval length follows a pattern regarding the gravida’s preceding pregnancy similar to the other early life indicators pattern, birth weight, placenta weight, gestation length and neonatal survival. Our results confirm and complete an immunological explanation of the indicators variations associated to the gravida’s preceding pregnancy.

## Introduction

A recent analysis of 27,243 neonates has shown a significant association between the mother’s preceding pregnancies and fetal development and neonatal survival [1]. A positive association was found when current conceptus and conceptuses of preceding pregnancies were of same sex, and a negative association when they were of opposite sex. An immunological hypothesis was put forward to explain the phenomenon. It is based on the sex-linked concepto-gravidic antigenic dissimilarity, due to paternal antigens of the conceptus, and it is capable of affecting fetal development and neonatal survival through a selective implantation process [1].

Early in gestation, the implantation of the blastocyst is a critical stage that a large number of conceptuses fail to achieve [2, 3, 4, 5, 6]. Implantation seems to be immunologically controlled [7, 8, 5]. Both, male and female conceptuses, can induce an immune reaction from the gravida. The concepto-gravidic dissimilarity, due to paternal antigens of the conceptus, favors the implantation of the blastocyst [9, 10, 11, 12, 5, 6, provides a selective advantage to the latter [13, 6, 14] and primes the gravida vis-à-vis some paternal antigens. Then, on the occasion of a subsequent pregnancy, the previous sensitization of the gravida reduces both the concepto-gravidic dissimilarity effect and the implantation advantage [15, 12, 16, 14]. Thus, the blastocyst carrying a given set of sex-linked paternal antigens, against which the mother has been previously sensitized, would implant less easily than the blastocyst whose mother has not been so exposed. However, if the former survives the implantation, it will be more successful than the latter in pursuing the gestation. This translates, in population terms, to survival of the strong and culling of the weak [13, 17, 18], as expressed in fetal development and neonatal survival differentials [1]. So, selective implantation favors in number blastocysts preceded by mother’s pregnancies of the opposite sex, and in strength, blastocysts preceded by mother’s pregnancies of same sex as their own. The first effect should be reflected in the post-implantation sex ratios, while the second should be shown in the development and survival differentials associated with the mother’s preceding pregnancies.

The present study investigates this phenomenon at a very early stage, that is during the interval separating the onset of the current pregnancy from the end of the immediately preceding pregnancy of the gravida. The influence of mother’s preceding pregnancies can be felt long before the onset of the current pregnancy. During the interpregnancy interval 70% of conceptions fail, including 30 before implantation, 30 after implantation but before the missed period, and 10 lost as miscarriage [19]. Therefore 60 % of the lost conceptuses are not clinically recognized. However they can delay the onset of a subsequent pregnancy by prolonging the interpregnancy interval [20]. Many factors are known to influence the IPI length. Our study investigated the variations of interpregnancy length associated with the mother’s preceding pregnancies.

## Materials and Methods

The basic information is derived from the Child Health and Development Studies (CHDS). Between August 1959 and September 1966, 20,000 pregnant women, members of the Kaiser Health Plan residing in the San Francisco - East Bay area, reported for prenatal care in some Kaiser clinics. They constituted the CHDS study population. Data on parents and children were obtained from interviews and medical records. The study population represented a broad range of economic, social and educational characteristics, and was not atypical of an employed population [21]. The current retrospective analysis is based on a sample of the CHDS population which included 7773 neonates who were alive at the time of hospital discharge, and whose mother had been interviewed about her reproductive history. Multiple births and newborns from diabetic mothers were excluded from the study.

The interval separating the beginning of the current pregnancy from the end of the immediately preceding pregnancy (IPI) was used as indicator of early undetected loss, which corresponds to a prolongation of this interval [20]. The current neonates were divided into cohorts according to their sex and the sex and number of the conceptuses of the mother’s preceding pregnancies (PP). Comparisons of IPI mean were made between pairs of cohorts of same sex and mother’s parity, but of different configuration of preceding pregnancies, mostly same sex preceding pregnancy (SSPP) and opposite sex preceding pregnancy (OSPP). The analysis included parity two, three and four.

## Results

### Characteristics of the participants in “the interpregnancy interval study”

**Table 1.**
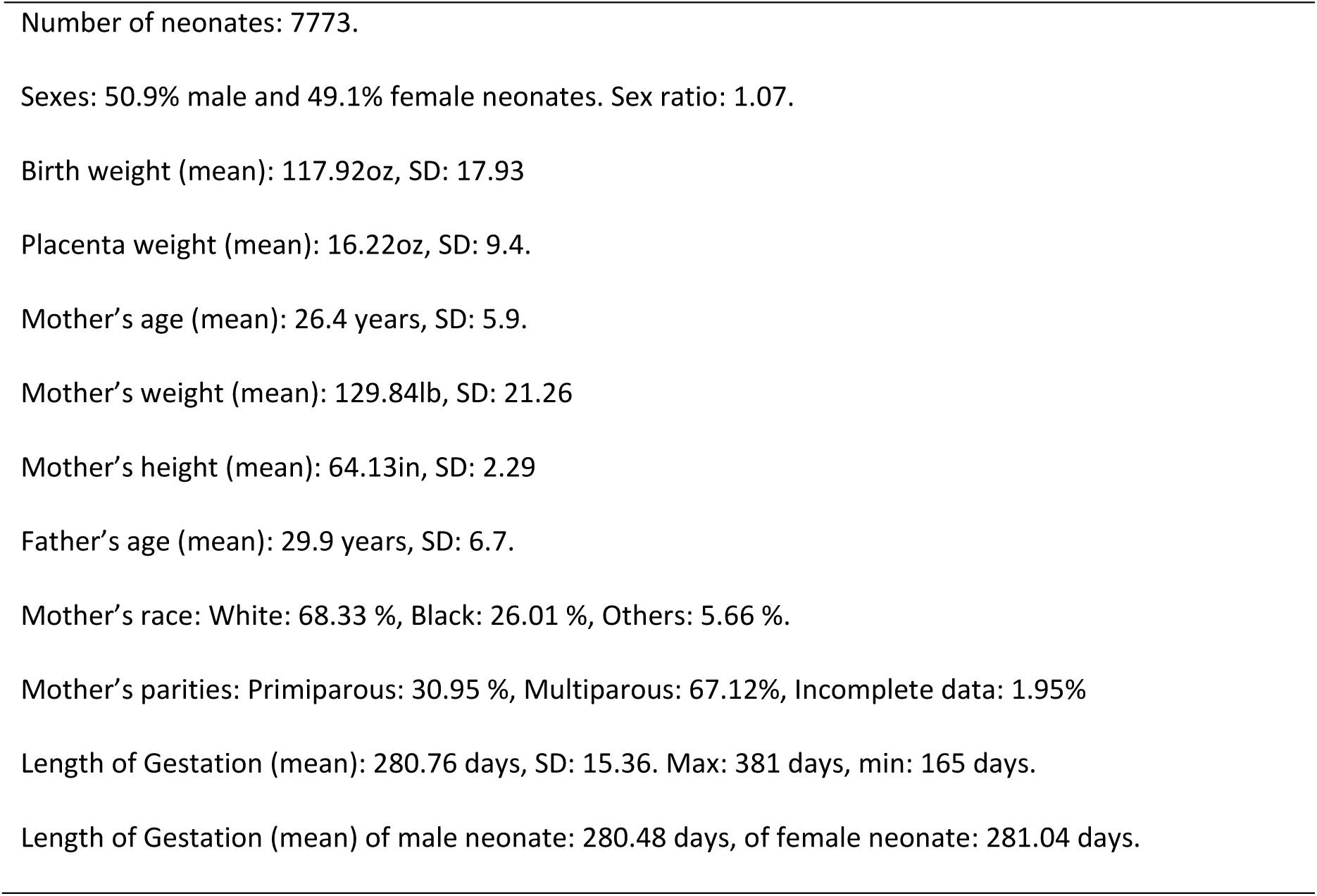
Characteristics of the participants in the Interpregnancy Interval Study.

### Interpregnancy Interval, sex of the current neonate and mother’s parity

In our study, the average length of time (IPI), separating the onset of the current pregnancy from the termination of the immediately preceding pregnancy was 779 days (26 months) for either female or male current pregnancy. It includes: 37% intervals of less than 1 year, 27% of between 1and 2 years, 26% of between 2 and 5 years, and 10% of 5 years or more. The interpregnancy interval of both male and female neonate increased with the parity of the mother (Table 2).

**Table 2.** Interpregnancy Interval (IPI), sex of the current neonate and mother’s parity.

| Male neonate IPI |  |  |  | Female neonate IPI |  |  |  |
| --- | --- | --- | --- | --- | --- | --- | --- |
| Parity | n | mean (days) | std dev | Parity | n | mean (days) | std dev |
| 2 | 911 | 701 | 857 | 2 | 926 | 713 | 869 |
| 3 | 719 | 835 | 909 | 3 | 711 | 839 | 952 |
| 4 | 474 | 887 | 909 | 4 | 429 | 843 | 879 |
| 5-7 | 531 | 741 |  | 5-7 | 516 | 762 |  |
| All | 2635 | 779 |  | All | 2582 | 779 |  |

### Interpregnancy interval and mother’s preceding pregnancy

At parity 2, (Table 3), the average interpregnancy interval (IPI) of a male neonate preceded by a male pregnancy of its mother was 635 days (21mo), and of a female neonate preceded by a male pregnancy was 787 days (26mo), a difference of 152 days (5mo): *p*=.0022 in favor of the latter. Likewise, the average IPI of a male neonate preceded by a female pregnancy of its mother was 834 days (28mo) and of a female neonate preceded by a female pregnancy was 687days (23mo), a difference of 147 days (5mo): *p*=.0156 in favor of the former. The average IPI of a male neonate preceded by a male pregnancy of its mother was 635 days (21mo), and preceded by a female pregnancy was 834 days (28mo), a difference of 199 days (6.6mo): *p*=.0005 in favor of the latter. Likewise, the average IPI of a female neonate preceded by a male pregnancy was 787 days (26 mo), and preceded by a female pregnancy was 687 days (23 months), a difference of 100 days (3.3 mo): *p* = .0484, in favor of the former.

**Table 3.**
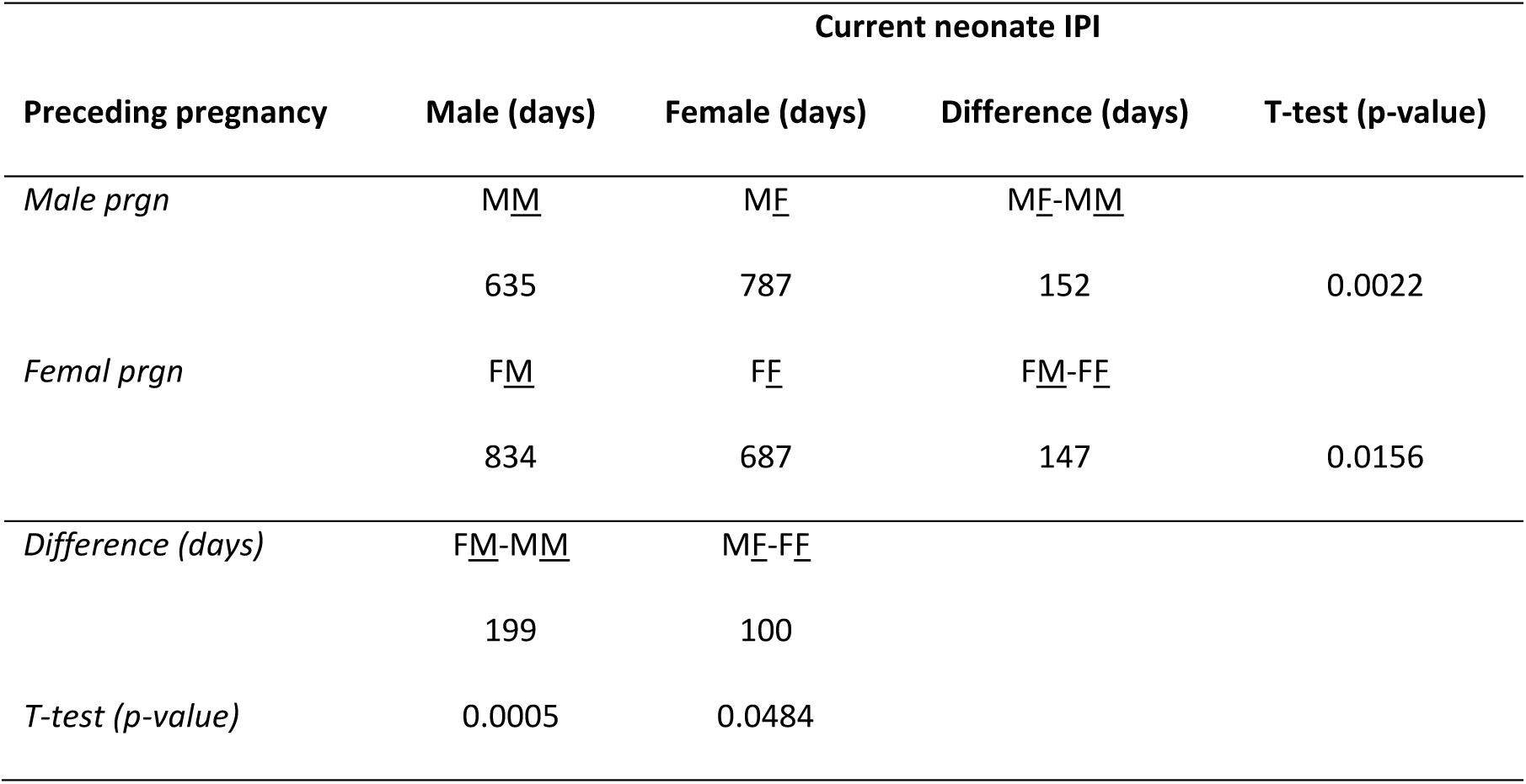
Interpregnancy interval (IPI) of the current neonate and mother’s preceding pregnancy, at parity 2.

At parity 3, (Table 4), the average interpregnancy interval (IPI) of a male neonate preceded by two male pregnancies of its mother was 876 days, and preceded by two female pregnancies was 903 days, a difference of 27 days in favor of the latter. Likewise, the average IPI of a female neonate preceded by two male pregnancies of its mother was 949 days and preceded by two female pregnancies was 926 days, a difference of 23 days in favor of the former.

Finally, at parity 4 the average IPI of a male neonate preceded by 3 male pregnancies of its mother was 948 days, and preceded by 3 female pregnancies was 903 days, a difference of 45 days in favor of the former. Likewise the average IPI of a female neonate preceded by 3 female pregnancies of its mother was 898 days, and preceded by 3 male pregnancies was 728 days, a difference of 170 days *p*= .0879, in favor of the former.

**Table 4.**
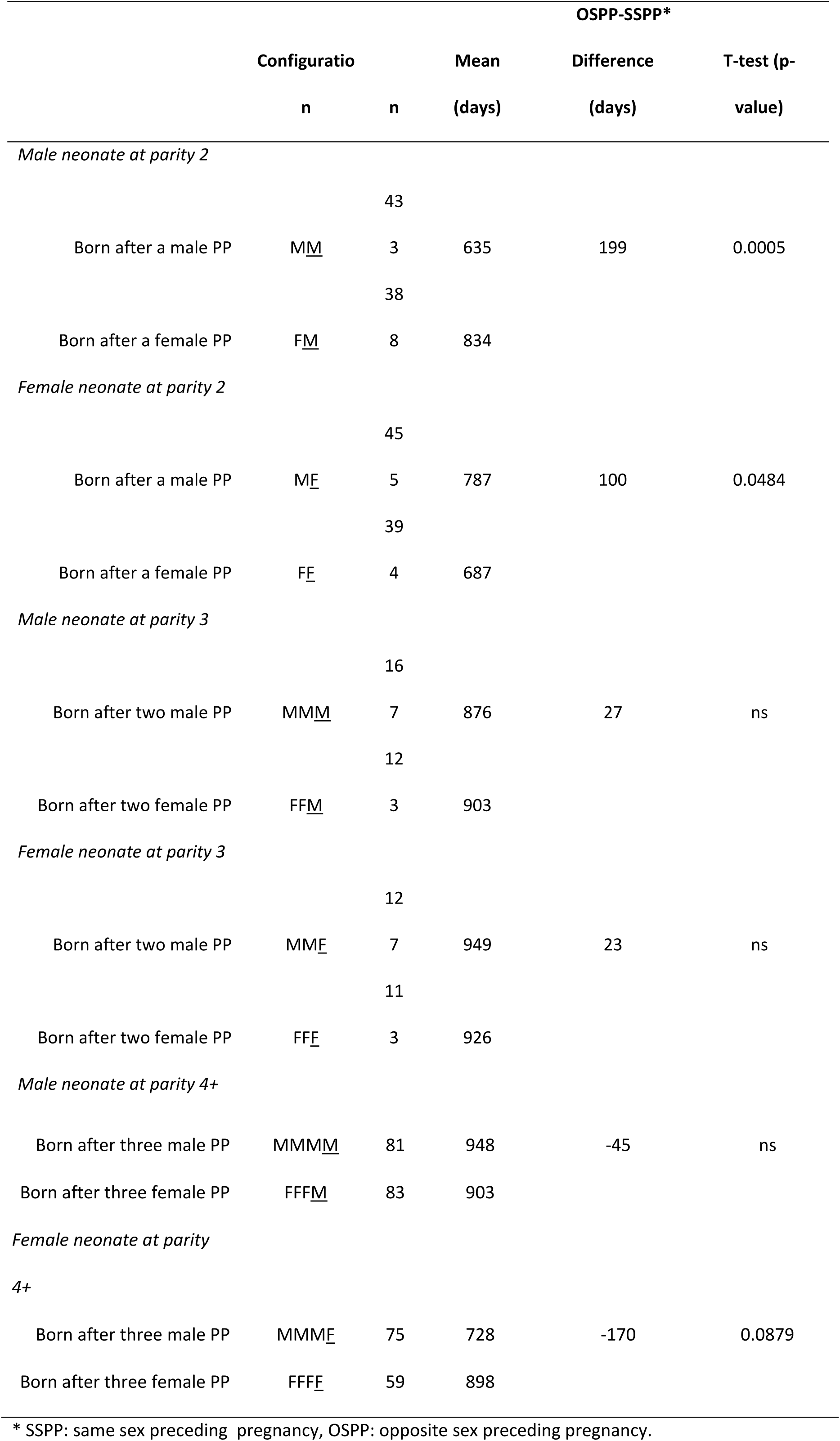
Interpregnancy Interval (IPI) of the current neonate and mother’s preceding pregnancies.

## Discussion

### Interpregnancy interval and mother’s preceding pregnancy

At parity 2, the interval between the termination of the preceding pregnancy and the onset of the current one is shorter if the conceptuses of the two pregnancies are of the same sex.

Thus, after a male preceding pregnancy, the IPI of the current male neonate is shorter than the IPI of the female neonate (635days vs 787days : *P=.0022*). And, after a female preceding pregnancy, the IPI of the current female neonate is shorter than the IPI of the male neonate (687days vs 834days : *P=.0156*). The difference in both cases is around 5 months, which is statistically significant (Table 3). In the same way, the IPI of a current male neonate born preceded by a male pregnancy is shorter than born preceded by a female pregnancy (635days vs 834days: *P=.0005*). And the IPI of a current female neonate born preceded by a female pregnancy is shorter than born preceded by a male pregnancy (687days vs 787days : *P=.0484*). The difference is respectively 6.6 mo for the male neonate and 3.3 mo for the female neonate, which are both statistically significant (Table 3).

At parity 3, the pattern of association between mother’s preceding pregnancy and current neonate’s interpregnancy interval persists at a lesser degree. Thus the difference between OSPP interval and SSPP interval is 27 days for the male neonate, and 23 days for the female neonate, and none of them is statistically significant (Table 4). Finally at parity 4+, the pattern of association reverses itself with SSPP interval longer than OSPP interval and negative difference between them. This latest reversal could be due to the small number of observations available for analysis. Or it could be due to the conceptus antigens loosing specificity with time or the response decreasing after a while [22, 23]. The value of the interpregnancy interval of the neonate (Table 5) increases between mother’s parity 2 and parity 3 of the neonates born after SSPP, as well as born after OSPP, initiating thus a limited parity-linked dose-effect.

**Table 5.**
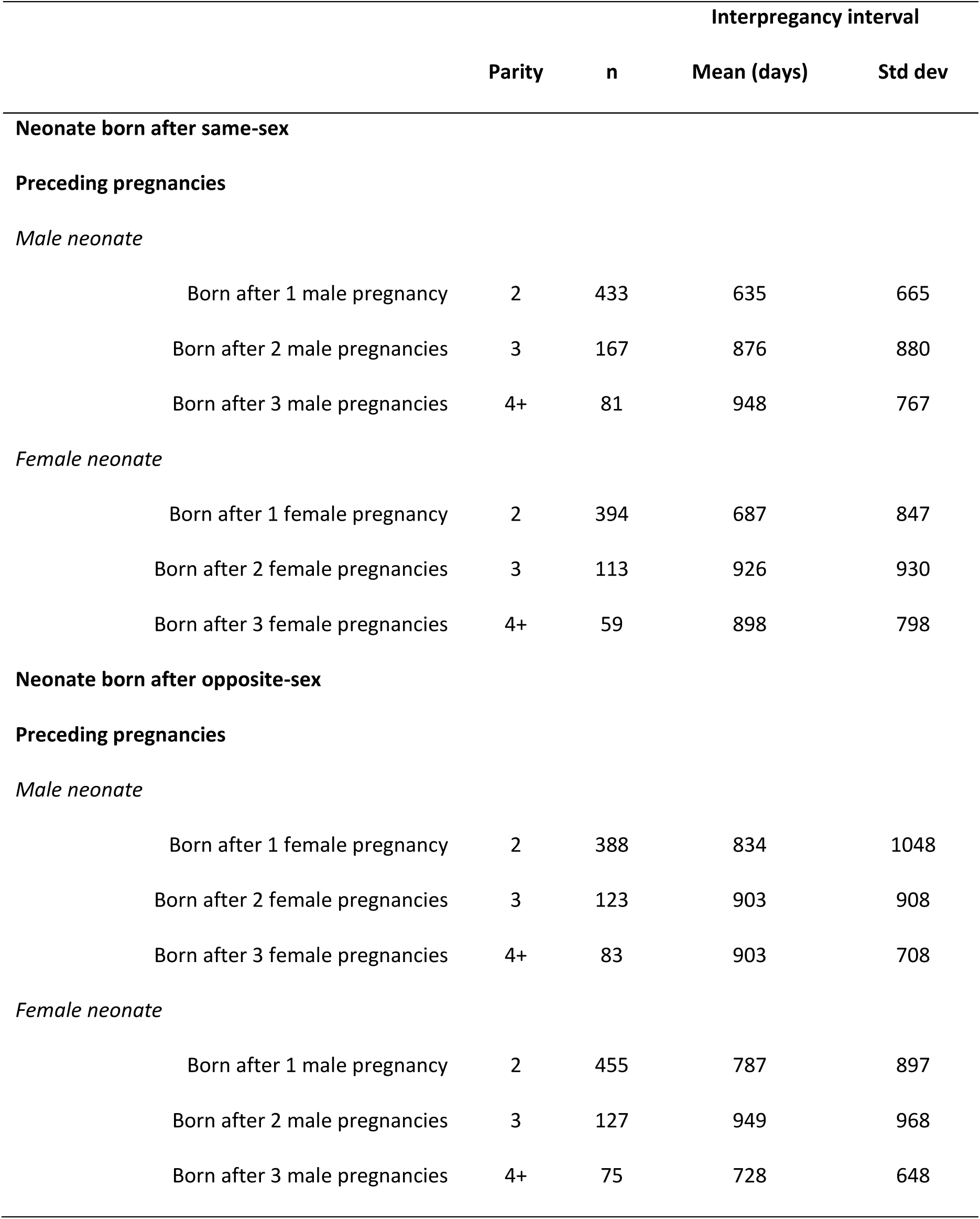
Interpregnancy Interval of the neonate born after same-sex preceding pregnancies (SSPP) or opposite-sex preceding pregnancies (OSPP).

### Immunological selective implantation and interpregnancy interval

A positive association was found between mother’s preceding pregnancy and interpregnancy interval when current pregnancy and preceding pregnancy were of the same sex, and a negative one when they are of opposite sexes. In a previous study [1], a similar pattern of association between current pregnancy and gravida’s preceding pregnancy was found for birth weight, placenta weight, gestation length and neonatal survival. An immunological hypothesis, based on the sex-linked concepto-gravidic antigenic dissimilarity due to paternal antigens of the conceptus was then proposed to explain the phenomenon. The selective implantation favors in number the blastocysts preceded by pregnancies of opposite sex, and favors in strength the blastocysts preceded by pregnancies of same sex as their own. Thus the blastocysts of multiparous gravidas would implant more easily after preceding pregnancies of opposite sex (OSPP) than after preceding pregnancies of same sex (SSPP). As a result, SSPP-blastocysts, ie MM and FF, would be fewer to successfully achieve implantation than OSPP-blastocysts, ie FM and MF. But the former, thanks to the selection process taking place at implantation, would be more successful than the latter in pursuing the gestation.

### Tentative explanatory mechanism

The influence of the gravida’s preceding pregnancy on the current one can be felt long before the onset of the latter. During the interpregnancy interval, 70% of all conceptions fail, including 30% before implantation, 30% after implantation but before the missed period, and 10% lost as miscarriage [19]. Therefore 60% of the lost conceptuses are not clinically recognized, and not counted as ‘preceding pregnancy’. However they can delay the onset of the current pregnancy by prolonging the IPI [20]. According to the immunological hypothesis, the blastocysts of the conceptions occurring during the interpregnancy interval would implant more easily after a preceding pregnancy of opposite sex (OSPP-blastocyst) than after a preceding pregnancy of same sex as its own (SSPP-blastocyst). In population terms, assuming a primary sex-ratio close to 1/1, more SSPP-conceptuses than OSPP- conceptuses would be eliminated before implantation while more OSPP-conceptuses than SSPP-conceptuses would be eliminated after implantation. Because conceptus loss before implantation has a shorter life than conceptus loss after implantation, a differential impact would result on IPI in relation to the preceding pregnancies. The IPI of the current neonate succeding to preceding pregnancies of same sex as its own would be shorter than the IPI of a current neonate succeding to preceding pregnancies of opposite sex. In our study:

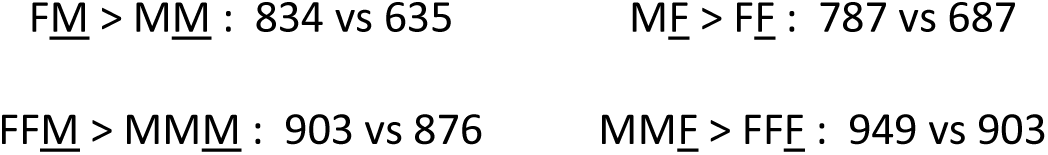

## Conclusion

Our study confirms and completes the role played by the preceding pregnancies of the gravida in the development and survival of her current conceptus. Specifically the study shows a statistically significant association between preceding pregnancies and interpregnancy interval. In order to test the consistency of a correlation between interpregnancy interval and preceding pregnancy a prospective approach with large samples would be necessary.

